# Latitudinal patterns in intertidal ecosystem structure in West Greenland suggest resilience to climate change

**DOI:** 10.1101/2021.01.05.419028

**Authors:** Jakob Thyrring, Susse Wegeberg, Martin E. Blicher, Dorte Krause-Jensen, Signe Høgslund, Birgit Olesen, Jozef Wiktor, Kim N. Mouritsen, Lloyd S. Peck, Mikael K. Sejr

## Abstract

Climate change has ecosystem-wide cascading effects. Little is known, however, about the resilience of Arctic marine ecosystems to environmental change. Here we quantify and compare large-scale patterns in rocky intertidal biomass, coverage and zonation in six regions along a north-south gradient of temperature and ice conditions in West Greenland (60–72°N). We related the level and variation in assemblage composition, biomass and coverage to latitudinal-scale environmental drivers. Across all latitudes, the intertidal assemblage was dominated by a core of stress-tolerant foundation species that constituted >95% of the biomass. Hence, canopy-forming macroalgae, represented by *Fucus distichus* subsp. *evanescens* and *F. vesiculosus* and, up to 69 °N, also *Ascophyllum nodosum*, together with *Semibalanus balanoides*, occupied >70% of the vertical tidal range in all regions. Thus, a similar functional assemblage composition occurred across regions, and no latitudinal depression was observed. The most conspicuous difference in species composition from south to north was that three common species (the macroalgae *Ascophyllum nodosum*, the amphipod *Gammarus setosus* and the gastropod *Littorina obtusata*) disappeared from the mid-intertidal, although at different latitudes. There were no significant relationships between assemblage metrics and air temperature or sea ice coverage as obtained from weather stations and satellites, respectively. Although the mean biomass decreased >50% from south to north, local biomass in excess of 10 000 g ww m^-2^ was found even at the northernmost site, demonstrating the patchiness of this habitat and the effect of small-scale variation in environmental characteristics. Hence, using the latitudinal gradient in a space-for-time substitution, our results suggest that while climate modification may lead to an overall increase in the intertidal biomass in north Greenland, it is unlikely to drive dramatic functional changes in ecosystem structure in the near future. Our dataset provides an important baseline for future studies to verify these predictions for Greenland’s intertidal zone.

## 1. Introduction

The rocky intertidal zone is one of the most studied marine habitats, and has provided a wealth of information about the ecological processes that shape assemblage structure and dynamics. The Arctic accounts for more than 30 % of the world’s coastline (Lantuit et al., 2012), but Arctic intertidal habitats have received little attention, and rocky Arctic shores were until recently thought to be sparsely colonized (Ellis, 1955; Węsławski, Wiktor, Zajaczkowski, & Swerpel, 1993). Scattered studies from different sub-Arctic and Arctic rocky shores on Svalbard (Kuklinski & Barnes, 2008; Węsławski, Dragańska-Deja, Legeżyńska, & Walczowski, 2018; Węsławski, Wiktor Jr., & Kotwicki, 2010), Iceland (Ingolfsson, 1992; Ingólfsson, 1996) and southern Greenland (Blicher et al., 2013; Høgslund et al., 2014; Ørberg et al., 2018) have documented species diversity, and revealed high biomasses locally, but over 95% of the Arctic intertidal zone remains unexplored. For instance, the physical parameters driving regional differences, and the rates of change in key ecosystem metrics, such as biomass and coverage, remain poorly understood, hindering assessments of climate change effects on the ecosystem. At lower latitudes, several studies have highlighted the intertidal zone as a harbinger of climate change impacts (Barry, Baxter, Sagarin, & Gilman, 1995; Helmuth et al., 2002), showing that increasing temperatures force intertidal species poleward (Pitt et al., 2010; Sanford *et al*., 2019), potentially causing ecosystem-wide effects as non-native foundation species establish, or natives disappear (Sorte et al., 2017). Although, the Arctic is warming at rates 2- to 3-fold greater than the global average (AMAP 2017), the scarcity of Arctic baseline and time-series data prevent quantifications of how this rapid warming affects intertidal organisms and ecosystem functioning.

Located in the middle of the North Atlantic, Greenland spans more than 20 degrees of latitude covering both sub-Arctic and Arctic ecosystems. In addition to the latitudinal gradient, the surrounding ocean currents create biogeographic differences between Greenland’s east and west coast. West Greenland is influenced by north-flowing warm water of Atlantic origin, while the East Greenland coast is dominated by cold south-flowing polar water from the central Arctic ocean via the East Greenland current. As a consequence, ocean temperatures, sea ice coverage and connectivity to source populations vary greatly around Greenland. Along the west coast, surface air temperatures decrease with latitude from an annual average of ∼0°C in the southwest to ∼-9°C in the north, but air temperatures have been steadily increasing since the 1990’s (Thyrring, Blicher, Sørensen, Wegeberg, & Sejr, 2017), resulting in declining sea ice coverage (Meredith et al., 2019). Although landfast patches of sea ice form within most fjords in the southwest, a large part of southwest Greenland is ice-free in winter, whereas winter ice cover is often established in and above the Disko Bay region (69°N), and seasonal ice-scour occurs following ice break-up at high latitudes (Gutt, 2001). In addition to sea ice, the Greenland Ice Sheet discharges large amounts of ice and meltwater into West Greenland fjords at rates which have increased 4-fold in recent decades in the southwest (Bevis et al., 2019), increasing the risk of the intertidal communities being exposed to ice scour. So, temperatures, sea ice coverage and ice scour are all changing latitudinally in Greenland, forming an environmental gradient along the coast. These central environmental factors are expected to contribute to large-scale alterations in Arctic coastal marine communities and productivity along the Greenland coast over space and time (Post et al., 2013; Krause-Jensen & Duarte, 2014).

Studying marine ecosystems along this gradient can provide an indication of what to expect at higher latitudes in a warmer future when conditions may be similar to current conditions in the south. Specifically, a quantification of communities and populations along the Greenland coast can provide knowledge on the relative importance of environmental drivers, as well as providing important baseline data for observing and quantifying future changes. For example, latitudinal studies from sub-tidal coastal habitats in Greenland have demonstrated distinct changes in population dynamics, often with depressed growth at increasing latitudes where ice cover is extensive (Blicher, Rysgaard, & Sejr, 2007; Krause-Jensen et al., 2012; Sejr, Blicher, & Rysgaard, 2009). For the intertidal realm, most studies have focused on the two common blue mussels species, *Mytilus edulis* and *M. trossulus* (Mathiesen *et al*., 2017); Blue mussels occur along the entire coastline, and while they can survive exposure to sub-zero air and water temperatures (Thyrring et al., 2015b, 2020a), they are confined to protective microhabitats such as, cracks and crevices (Blicher et al., 2013), and their abundance and vertical distribution decrease with latitude (Thyrring et al., 2017). However, quantitative studies of intertidal assemblage structure in Greenland are limited to a few sub-Arctic sites (Høgslund et al., 2014; Ørberg et al., 2018).

The aim of the present study was to increase knowledge on Arctic rocky shore assemblage structure and resilience. We quantified latitudinal differences in abundance and biomass of key intertidal species along the West Greenland coast between 60°N and 72°N, and tested the hypotheses that (1) assemblage composition change and biomass and coverage decline towards higher (more Arctic) latitudes, and (2) intertidal species is progressively limited to lower intertidal heights at higher latitudes in the intertidal zone at high latitudes in response to increasing environmental stress.

## 2. Material and Methods

### 2.1 Field sampling

Local environmental conditions affect biomass and communities across small spatial scales. To reduce confounding effects of local-scale processes (e.g. within fjords) on large-scale climate related effects (Archambault & Bourget, 1996; Høgslund et al., 2014), we sampled a total of 320 plots distributed among 56 rocky intertidal sites in six different regions in West Greenland (Fig. 1). Sampling was conducted in July/August over the years 2011-2013 (Table 1). The aim was to sample enough sites within each region to capture local variability, which is necessary for an analysis aimed at identifying large scale changes. Thus, the sites were selected to a) represent a range of wave exposure, surface orientation and surface heterogeneity within each region, and b) ensure comparability in those conditions between regions. Sampling was conducted in the mid-intertidal during low tide at transects parallel to the shoreline using seven replicate plots of 0.0625m^2^ (25 ×25 cm frames) at each site following the sampling protocol of Høgslund et al. (2014). Within each frame, the total coverage of the intertidal assemblage, including the overhanging canopy from macroalgae attached outside the frame, was visually assessed. After harvest of the canopy-forming algae attached within the frame, the understory coverage of barnacles (*Semibalanus balanoides*) was equally assessed. All macroalgal- and macrozoobenthic species (larger than 0.5 cm) occurring within the frame, were subsequently collected, sorted, counted (fauna species) and weighed, except for barnacles (*Semibalanus balanoides*), for which only coverage was estimated. For all sites, the mid-intertidal was defined as half the maximum tidal amplitude, which was calculated from the Greenland tidal table (obtained from ocean.dmi.dk). The height on the shore was measured following Høgslund et al. (2014). At the northernmost region (Upernavik, Fig. 1) sampling was expanded to also include a lower intertidal level; and seven additional replicates were sampled 30 cm below the mid-intertidal (Table 1). In all regions and at all sampling sites, the vertical distribution of species was additionally registered along three vertical transect lines extending from the upper distribution limit of intertidal macroalgae to the lowest low water line. For every 25 cm along each vertical transect (starting from the lower distribution limit), the occurrence of species was recorded along a 20 cm horizontal line. At the two southern most locations only macroalgae (Cape Farewell) or canopy forming macroalgae (Ydre Kitsissut) were registered in vertical transects. The corresponding tidal heights were determined as described in Høgslund et al. (2014).

**Table 1:**
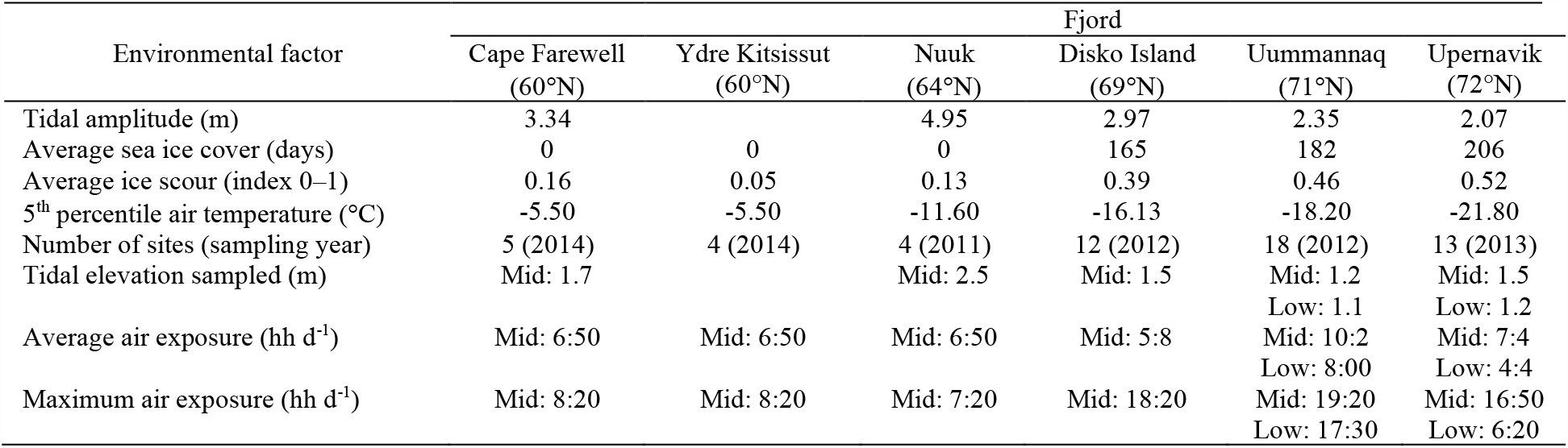
Environmental characteristics of the six study regions in West Greenland.

**Figure 1:**
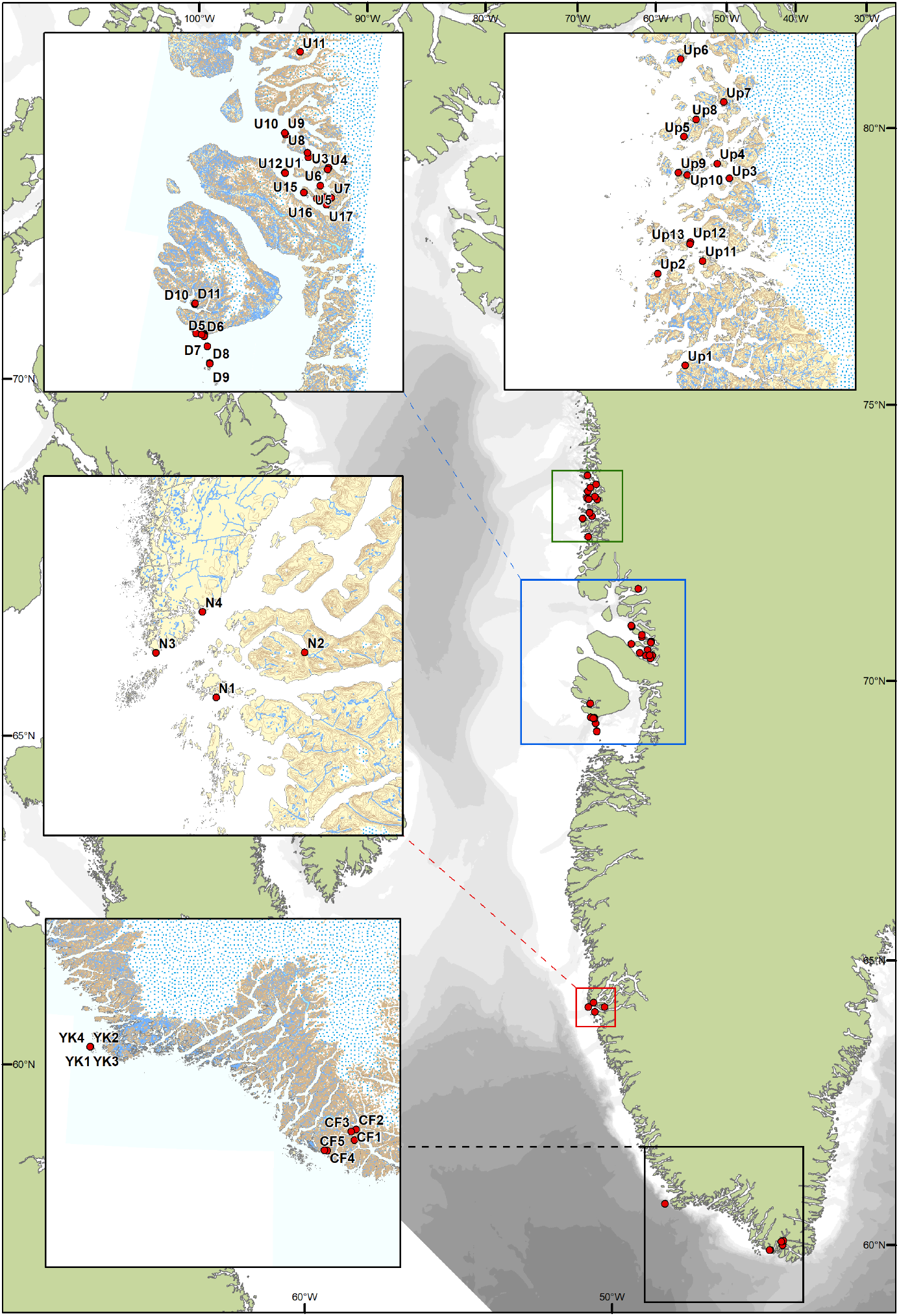
Position of 56 sampling sites nested within six regions in West Greenland: Cape Farewell (sites CF1–CF5), Ydre Kitssissut (sites YK1–YK4), Nuuk (sites N1–N4), Disko Island (sites D1– D12), Uummannaq (sites U1–U18), and Upernavik (sites Up1–Up13).

Macroalgae species from the frame samples were identified based on Pedersen (2011) to lowest possible taxonomic level using a stereomicroscope. Coverage is presented as 1) total coverage measured as percentage of frame covered before removal of the canopy macroalgae, and 2) understory *S. balanoides* coverage measured after removal of macroalgae. Biomass (wet weight, ww) only included species attached inside the frame and was measured in the lab and rounded to the nearest 0.001 g. Macrozoobenthic species were sorted and identified to lowest taxonomic level (Hayward & Ryland 1995), abundances were expressed as number of individuals, and biomass (ww) rounded to the nearest 0.001 g.

### 2.2 Environmental characteristics

Four environmental parameters were quantified for each region; length of exposure time to air during low tide, air temperature, sea ice coverage and ice scour. For each sampling locality an annual average and maximum air exposure time per tidal height was calculated based on 2013-tidal models (10 min resolution) for the nearest settlements, obtained from the Danish Meteorological Institute (*Pers. com*. Palle Bo Nielsen, Center for Ocean and Ice). Low air temperature stress can be important in the Arctic region, and the air temperatures were calculated as the fifth^th^ percentile averaged over a ten-year period (year 2007 to 2016) from data derived from local weather stations in Qaqortoq (station no 4286) Nuuk (station no 4250); Disko Island (station no 4220/4224); Uummannaq (station no 4212/4213) and Upernavik (station no 4210/4211) (Cappelen, 2017). The variation in the sea ice coverage period was characterized as a five-year mean based on visible-band images from the MODIS instrument on NASA’s Terra satellite. Indications of ice scour were assessed for each replicate plot, resulting in an ice scour index of 0 for no ice scour and 1 for indications of ice scour. These indices could only be obtained if vegetation were present within the replicate plots as it is based on deformities of *Fucus* spp. If deformities were registered, it was presumed to be caused by ice scouring and the replicate plot was assigned 1. Deformities registered ranged from depauperated and /or damaged thallus, including thallus previous being damaged and from where regrowth is initiated, to complete destruction of plants only leaving reduced and crippled plants within crevices. Analyses of ice scour indications were performed from photos taken before sampling from each replicate plot.

### 2.3 Statistical analysis

Assemblage composition patterns between regions were depicted through ordination by nonmetric Multidimentional Scaling (nMDS) based on biomass data processed through the *vegan* package in R using a Bray-Curtis dissimilarity metric (Oksanen et al., 2018; R Core, 2019). Generalized linear mixed effect models (GLMM; *lme4* R package (Bates et al., 2015)) were used to test for significant effects of latitude on coverage (Binomial distribution on percentage data) and biomass (Gaussian distribution on continuous data) among regions using within-region sampling sites as a random intercept, as this accounted for dependency among sites located within a region (Zuur et al., 2009; Zuur & Ieno, 2016). Length of exposure time to air during low tide, fifth percentile air temperature, sea ice coverage and ice scour were included as explanatory variables in the full models, and all model assumptions were verified by plotting residuals versus fitted values (Zuur, Ieno, & Elphick, 2010; Zuur & Ieno, 2016). Following data exploration, biomass was log-transformed, and total coverage and *S. balanoides* coverage was square root-transformed, to avoid overdispersion and to ensure model assumptions were met. Estimated marginal means (EMM) were used to depict the significant effects of regions, using the *emmeans* R package (Searle et al., 1980; Lenth, 2019). Generalized linear models (GLM), were used to test for vertical differences in average number of species (Poisson distribution), algal biomass (Gaussian distribution), total coverage (Quasibinomial distribution) and *S. balanoides* coverage (Quasibinomial distribution) in the northernmost regions where sampling represented both mid-intertidal and lower-intertidal levels. Data were presented using the R packages *ggplot2* (Wickham, 2016) and *lattice* (Deepayan, 2008), and standard deviation (SD).

## 3. Results

### 3.1 Environmental characteristics

Tidal dynamics along the Greenland coast change from semidiurnal cycles with 3 to 5 m maximum amplitude at 60–64°N, to mixed tidal cycles and 2 to 3m amplitude further north (Table 1). Thus, the average and maximum air exposure time during low tide varied among regions. The longest exposure time of 19.4 h per diurnal cycle in the mid-intertidal zone was recorded in Uummannaq (71°N) and the shortest (7.4 h) in Nuuk (64°N; Table 1). Average fifth percentile air temperatures across years decreased from −5.5°C in Cape Farewell (60°N) to −21.8°C in Upernavik (72°N), and the length of the sea ice-covered period increased latitudinally with south Greenland regions (Cape Farewell, Ydre Kitssissut (60°N), Nuuk) being ice-free year-round, to an ice-covered period of 206 days year^-1^ in the northernmost region, Upernavik (72°N; Table 1). Average ice scour increased latitudinally from 0.05 ± 0.23 at Ydre Kitssissut to 0.53 ± 0.50 at Upernavik (Table 1), while glacial ice was observed in all regions, and drift ice from the east coast of Greenland was observed at Cape Farewell.

### 3.2 Assemblage composition

We registered a total of 53 species in the mid-intertidal zone across six regions and 56 sites, distributed between 29 macroalgal species and 24 macrozoobenthic species. The macroalgal species were represented by 13 Chlorophyta, 11 Ochrophyta and 5 Rhodophyta, and the macrozoobenthic species by 10 Mollusca, 7 Arthropoda, 3 Cnidaria, 2 Annelida and 2 Platyhelminthes (Table 2). The nMDS ordination based on biomass (2-dimension, final stress = 0.118) illustrated high similarity of species composition among the six regions (Fig. 2). The most conspicuous difference in species composition from south to north, was that three common species disappeared completely from the mid-intertidal zone along the latitude gradient: The amphipod *Gammarus setosus* was only found in Cape Farewell (60°N), while *G. oceanicus* was found in all regions. The gastropod *Littorina obtusata*, otherwise common, was absent in Upernavik (72°N), and no mid-intertidal individuals of *Ascophyllum nodosum* were recorded in the biomass samples north of Nuuk (Table 2), although this species was also recorded lower in the tidal zone at Disko Island (Figure 6).

**Table 2:**
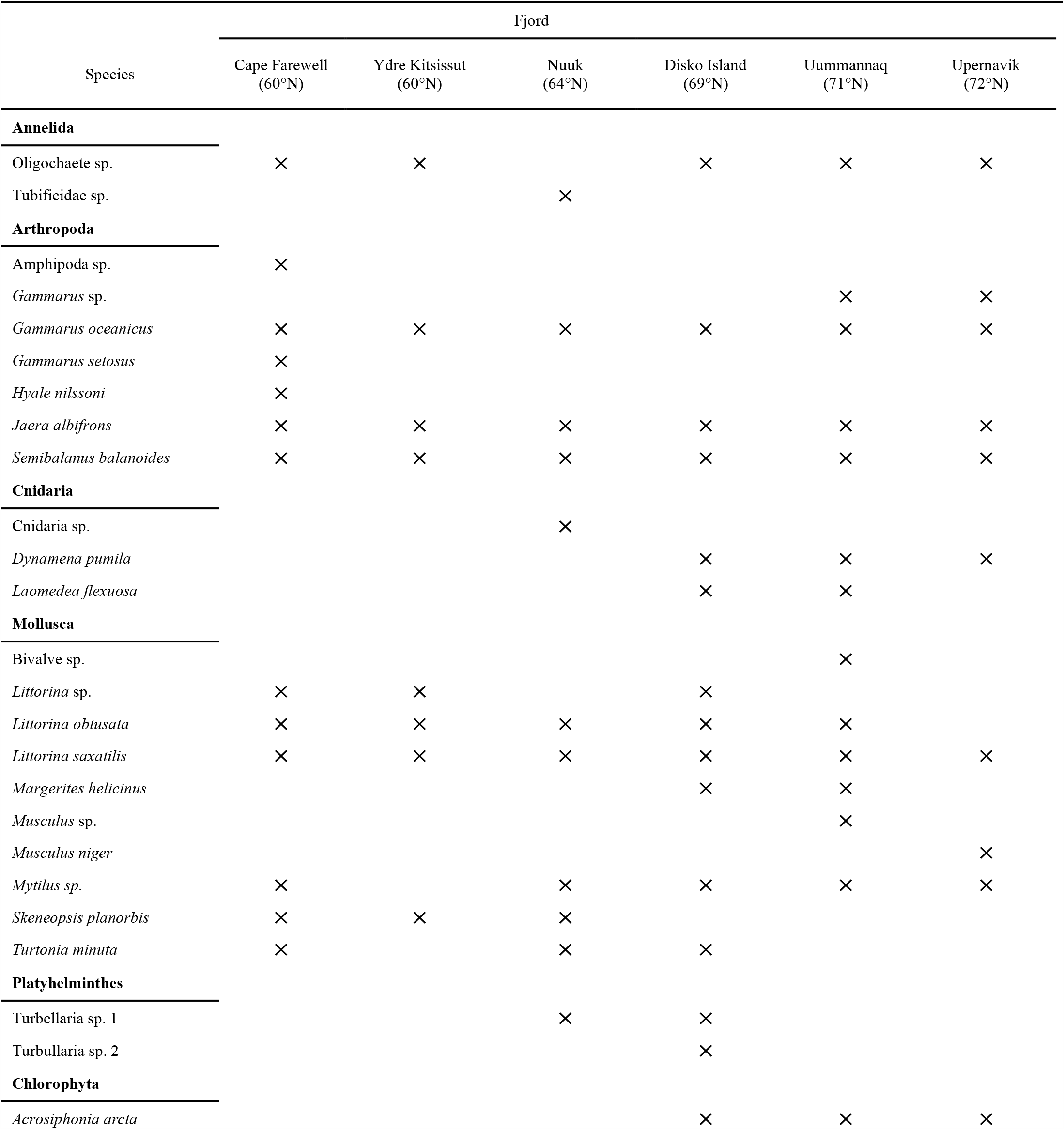
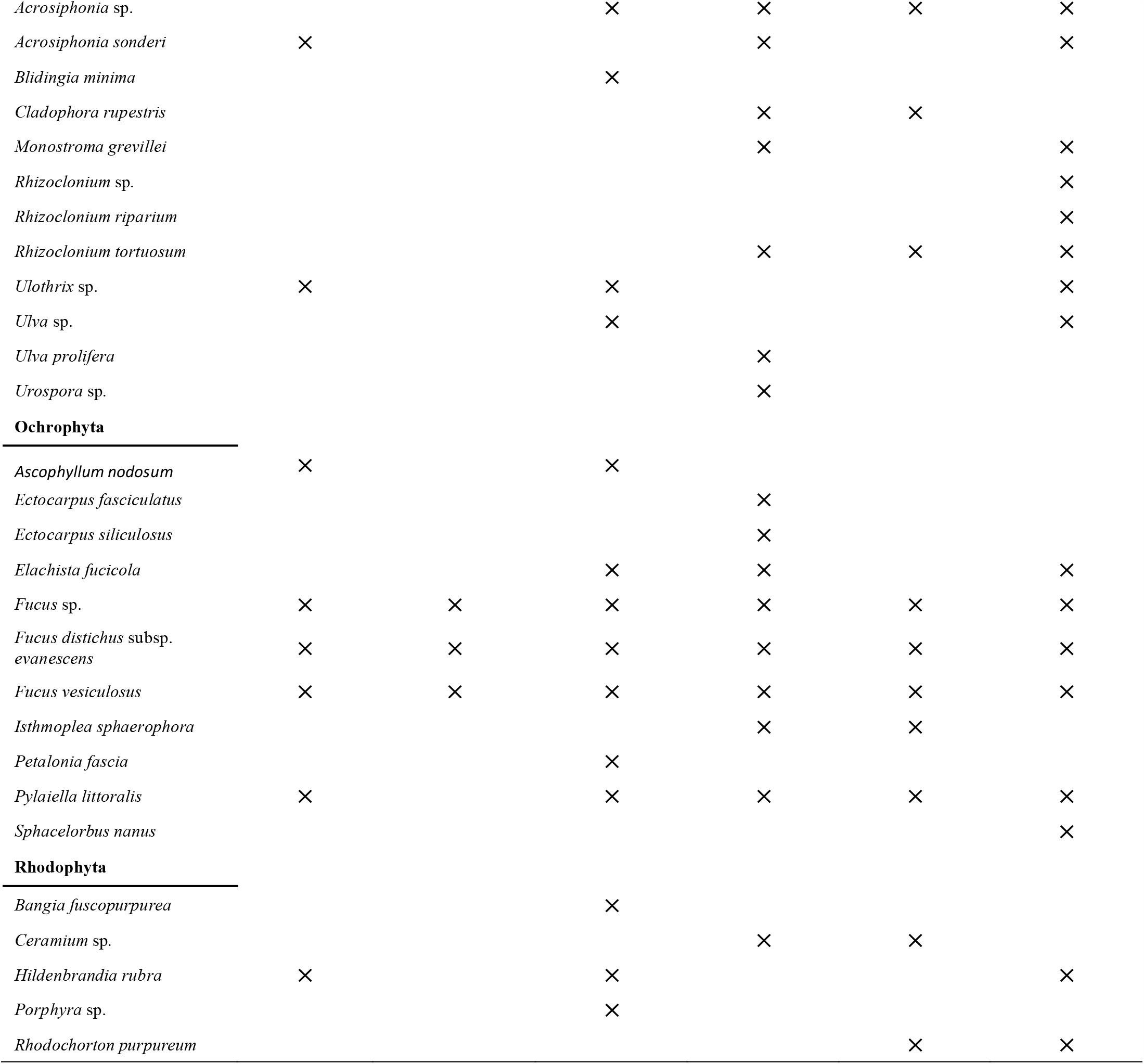
Species found in the mid-intertidal at the six studied regions in West Greenland. Please note, that for Ydre Kitsissut, only the fucoid foundation species were sampled together with the associated macrozoobenthic species.

**Figure 2:**
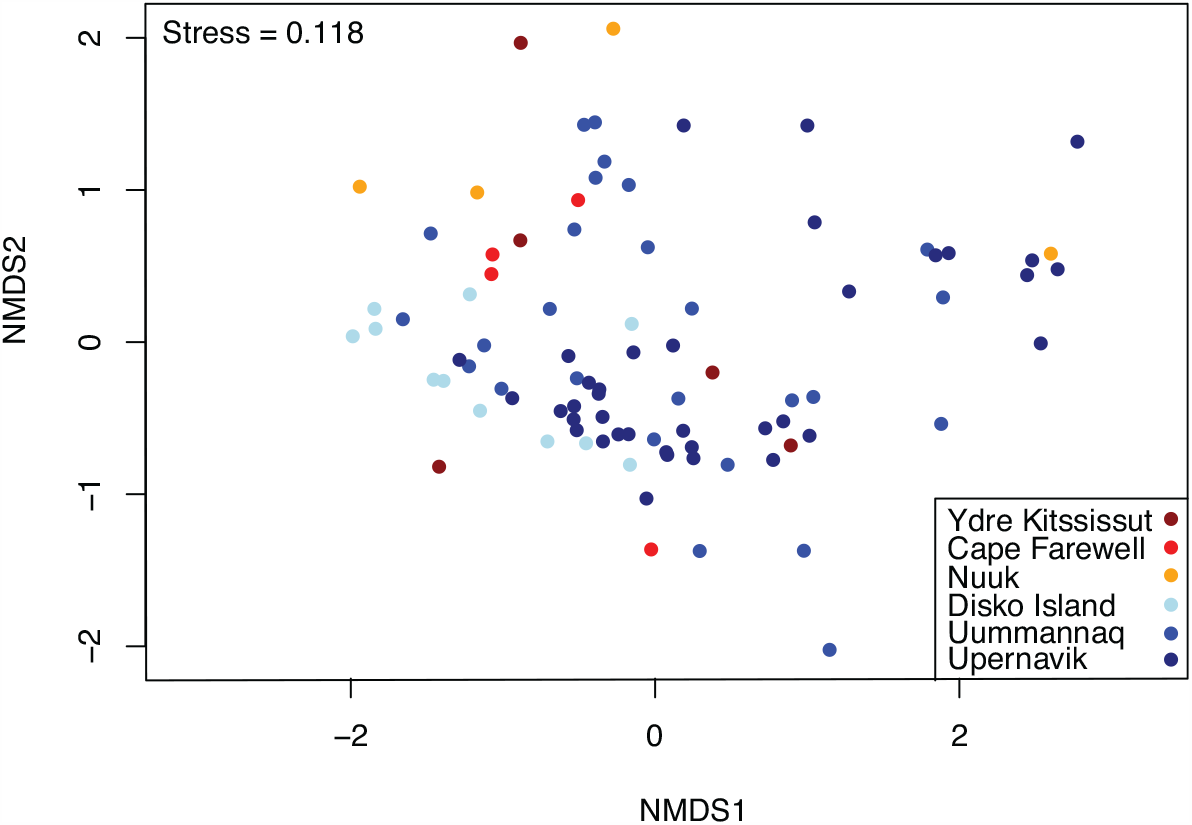
Representation of the species composition of the mid-intertidal assemblage along Greenland’s west coast from 60 to 72°N visualized through a 2-dimensional nonmetric multidimensional scaling (nMDS) ordination.

**Figure 3:**
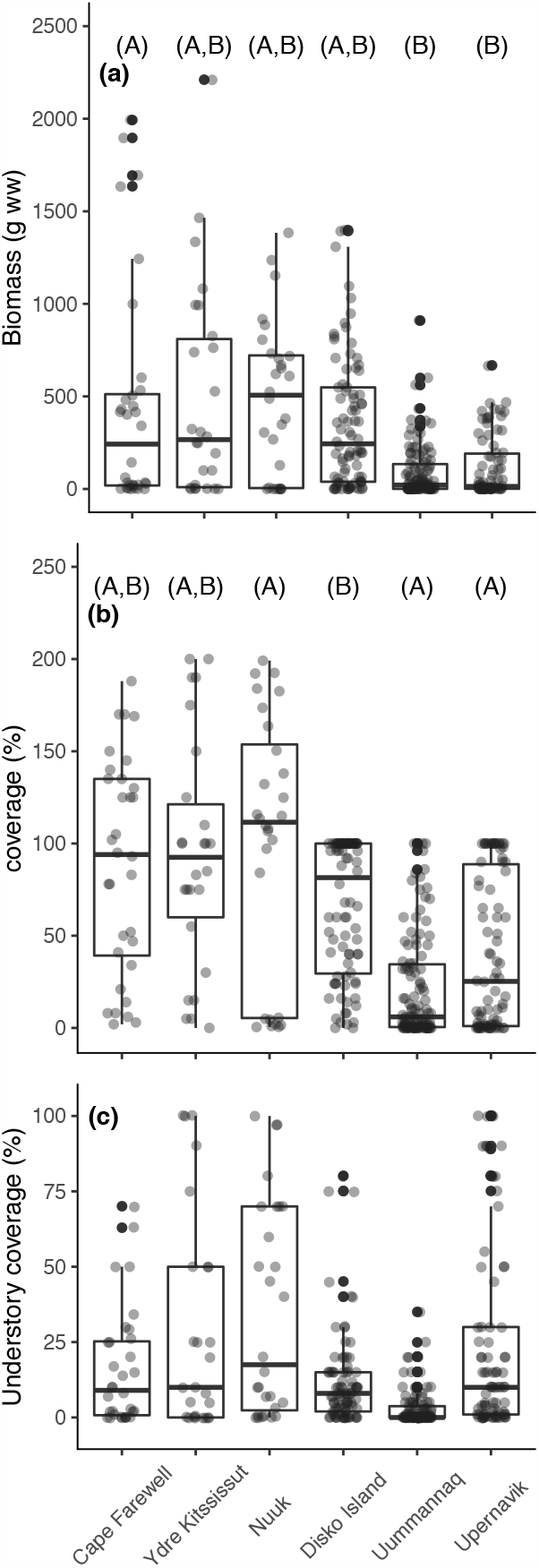
Boxplots of species biomass (g ww) (a), total coverage (%) (b) and understory *Semibalanus balanoides* coverage (%) after removal of macroalgae (c) per frame (0.0625m^2^) in six West Greenland regions from 60 to 72°N. The horizontal line in each boxplot is the median, the boxes define the hinges (25%–75% quartile) and the whisker is 1.5 times the hinges. Black dots represent data outside this range. Letters above boxplots indicate pairwise significance; groups with the same letter do not differ significantly (P < 0.05) among regions.

**Figure 4.**
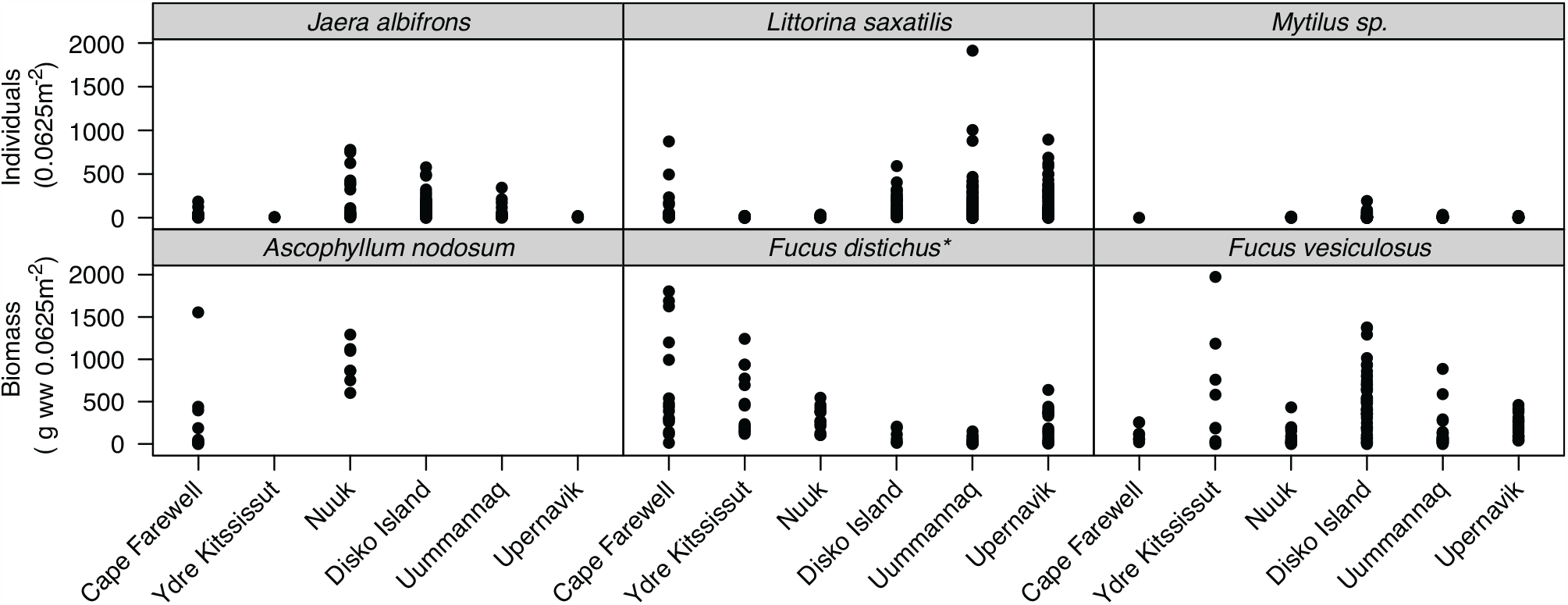
Mid-intertidal biomasses and abundances of the six most common species in six West Greenland regions from 60 to 72°N. Abundance (Individuals 0.0625m^-2^): *Jaera albifrons, Littorina saxatilis, Mytilus* sp. Biomass (g ww 0.0625m^-2^).: *Ascophyllum nodosum, Fucus distichus* (*Full name *Fucus distichus* subsp. *evanescens*) and *Fucus vesiculosus*.

**Figure 5:**
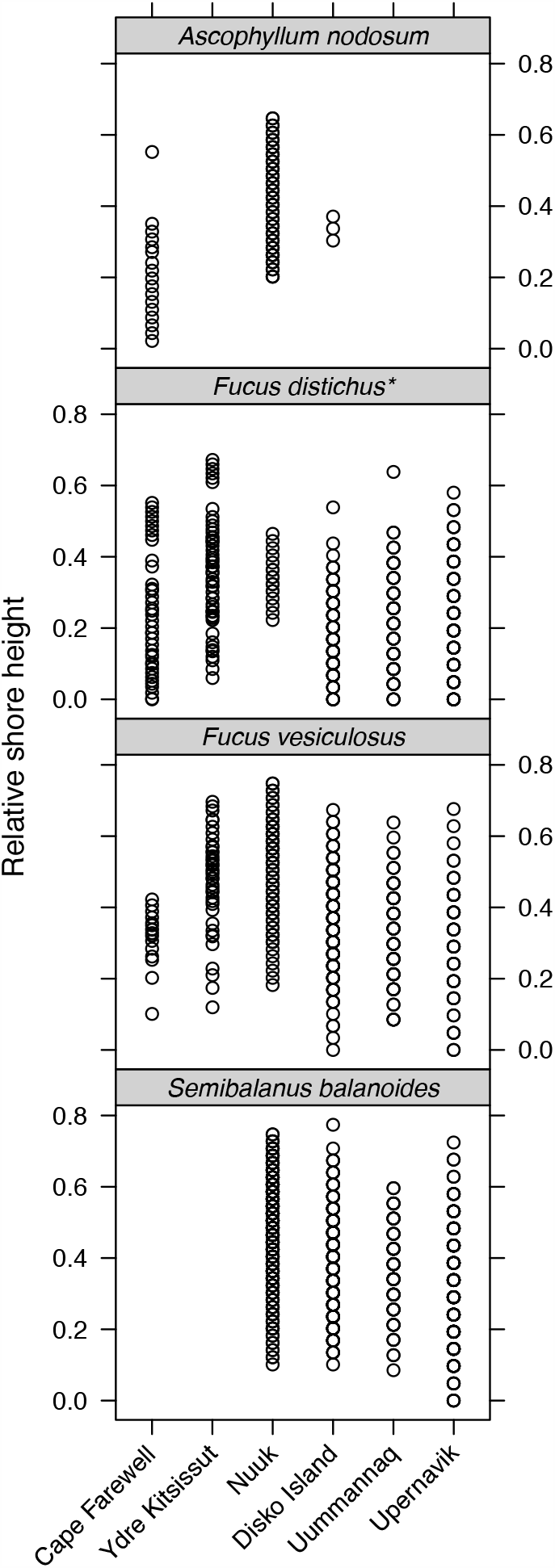
The vertical zonation (relative to max local tidal amplitude) of the four most common species in the intertidal zone of six West Greenland regions from 60 to 72°N. At the two southern most locations (Cape Farewell and Ydre Kitsissut) only macroalgal species were registered at vertical transects. *Full name *Fucus distichus* subsp. *evanescens*)

**Figure 6:**
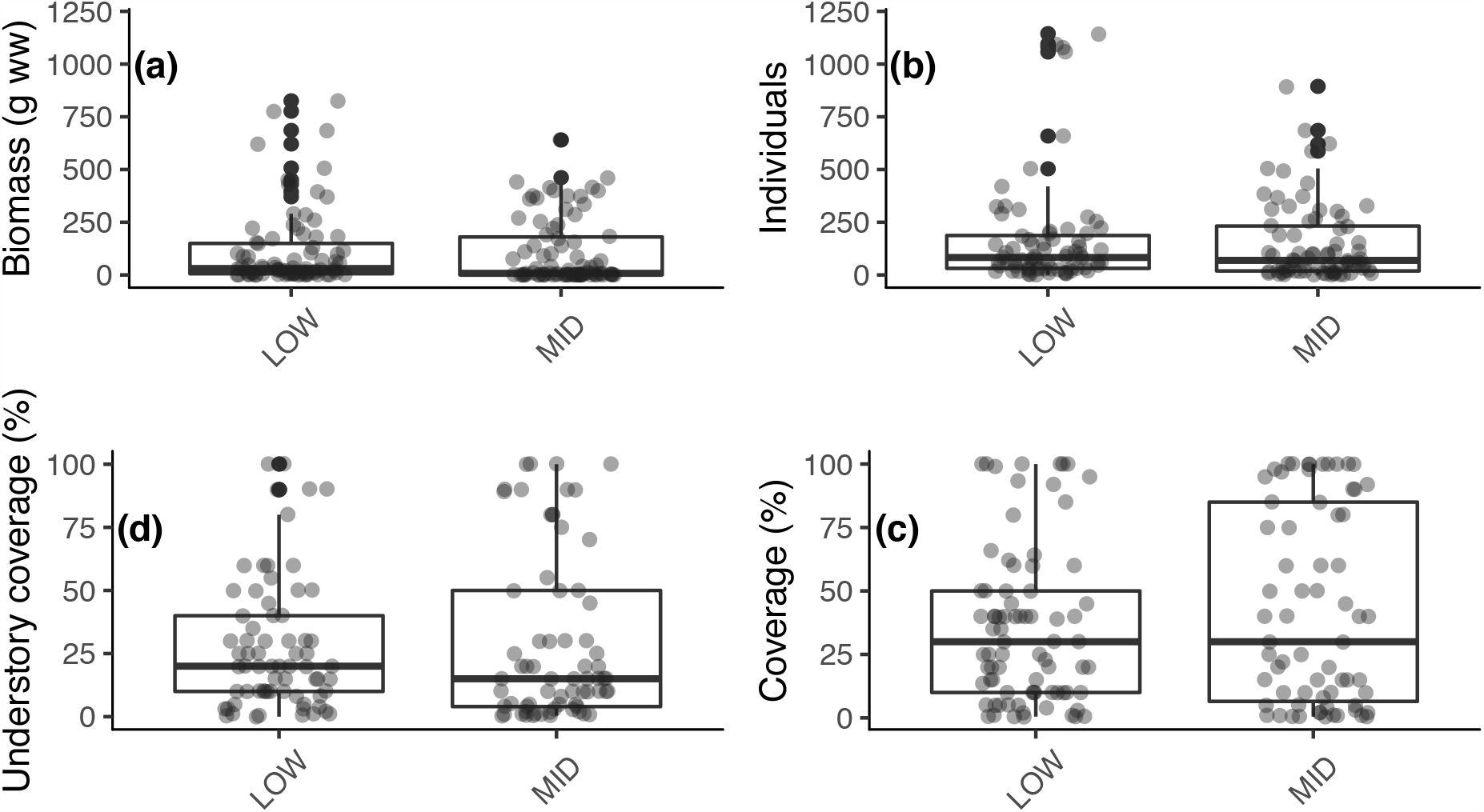
Total biomass (g ww) (a), abundance of benthic species (b), total coverage (c) and understory *Semibalanus balanoides* coverage (d) 0.0625m^-2^ at the mid-intertidal (MID) and 30 cm below the mid-intertidal (LOW) in Upernavik, West Greenland (72 °N).

### 3.3 Abundance, biomass and coverage

Total biomass ranged from 0–2209.9 g ww 0.0625m^-2^ in South Greenland at Ydre Kitssissut, and 0– 665.2 g ww 0.0625m^-2^ in the Upernavik area (Fig. 3a). There was a significant change in biomass along the latitudinal gradient (GLMM: p < 0.0001, Table 3), with a significantly lower biomass at Uummannaq (EMM: p = 0.0001) and Upernavik (EMM: p = 0.0004) than Cape Farewell (Fig 3a). This decrease was not significantly correlated to longer duration of ice coverage (GLMM: p = 0.38), maximum exposure time to air (GLMM: p = 0.38), air temperatures (GLMM: p = 0.73) or ice scour (GLMM: p = 0.19) (Table 3). In the mid-intertidal, canopy-forming brown macroalgal species constituted >95% of the biomass. This functional group was dominated by *A. nodosum, Fucus vesiculosus* and *F. distichus* subsp. *evanescens* (Supplementary Fig. 1). *A. nodosum* was found in Cape Farewell and Nuuk with biomasses exceeding 1500 g ww 0.0625m^-2^ and 1300 g ww 0.0625m^-2^, respectively (Fig. 4). Mid-intertidal *Fucus* spp. were found in all six regions; the highest biomass of *F. distichus* subsp. *evanescens* was recorded in southern Greenland, while *F. vesiculosus* was most abundant at Ydre Kitssissut and at Disko Island (Fig. 4). The three most abundant macrozoobenthic species were the isopod *Jaera albifrons*, the blue mussel *Mytilus* sp., and the grazer *Littorina saxatilis*, which occurred in densities of 0–1910 individuals 0.0625m^-2^ (Fig. 4).

**Table 3.** Generalized linear Mixed Effects model results for the best models with biomass or total coverage of the intertidal assemblage in six regions along Greenland’s west coast (60 to 72°N) as response variable, air temperature (fifth^th^ percentile), maximum air exposure, sea ice cover and ice scour as fixed effects, and sites nested within the six regions as random effects.

| Response variable | Coefficients fixed effects |  |  |  | Variance random effects |
| --- | --- | --- | --- | --- | --- |
|  | 5<sup>th</sup> percentile air temperature | Maximum air exposure | Sea ice cover | Ice scour |  |
| Biomass | -0.03, P = 0.73 | -0.19, P = 0.38 | -0.19, P = 0.38 | 0.29, P = 0.19 | 0.0583 |
| Total coverages | -0.03, P = 0.51 | 0.007, P = 0.95 | 0.005, P = 0.58 | -0.07, P = 0.65 | 2.84e-15 |

Total coverage (macroalgae and understory *S. balanoides* coverage combined) varied among regions (GLMM: p = 0.003) with a significantly higher coverage at Disko Island compared to Nuuk (MME: p = 0.02), Uummannaq (MME: p = 0.003) and Upernavik (MME: p = 0.023). The average total coverage showed a clear decline north of Disko Island from 67.2 ± 35.3% at Disko Island to 28.9 ± 30.1 % at Uummannaq (Fig. 3b), but neither ice coverage (GLMM: p = 0.58), maximum exposure time to air (GLMM: p = 0.95), air temperatures (GLMM: p =0.51) or ice scour (GLMM: p =0.65) correlated significantly with total coverage (Table 3). *S. balanoides* coverage displayed no significant latitudinal pattern with a local 100% coverage of the rock surface in Ydre Kitssissut, Nuuk and Upernavik (Fig. 3c).

### 3.4 Vertical distribution patterns

The recording of species along vertical transect lines extending from the lowest low water-tide line to the upper distribution limit of intertidal macroalgae, showed that the macroalgal vegetation belt occupied ∼70% of the entire tidal amplitude across all regions. This reflecte that the dominant macroalgal species (Fig. 5; *F. vesiculosus* and *F. distichus* subsp. *evanescens* did not change their vertical distributions across latitudes (Fig. 5; A full list of species occurring along the vertical transect is available in Supplementary Table 1). Likewise, *S. balanoides* inhabited ∼70–80% of the tidal amplitude in all regions where records were obtained. By contrast, the upper distribution limit for *A. nodosum* declined at Disko Island, relative to regions further south (Fig. 5). At Upernavik, where an extra set of plots was sampled at 30 cm below mid-intertidal, there were no significant effects of tidal height on biomass (GLM: p= 0.07), abundance of macrozoobenthic species (GLM: p= 0.48), total coverage (GLM: p= 0.20) or understory coverage (GLM: p= 0.46) (Fig. 6).

## 4. Discussion

We quantified the mid-intertidal assemblage in West Greenland along a latitudinal gradient from 60°N to 72°N. North of the Disko Island (69°N), the average biomass was >50% lower than in the southernmost regions. However, besides the absence of the foundation species *Ascophyllum nodosum* in north Greenland, the mid-intertidal assemblage dominance was maintained by the few habitat forming species (i.e. *Fucus vesiculosus, F. distichus* subsp. *evanescens* and *Semibalanus balanoides*) across all latitudes. The conspicuous poleward decline in biomass is consistent with studies on the terrestrial vegetation in West Greenland, which identify a transition zone from sub-Arctic to Arctic vegetation through the northern region of Disko Island (Fredskild, 1996). Similar patterns have also been reported for species residing in the subtidal around Greenland. Here the length of sea ice cover explains up to 47% of the variation in kelp production (Krause-Jensen et al., 2012), and the length of the productive open-water period explains more than 80% of growth differences among populations of the sea urchin *Strongylocentrotus droebachiensis* (Blicher et al., 2007). Hence latitudinal patterns of poleward decline in biomass of terrestrial and subtidal ecosystems in Greenland parallel to some extent the results reported here for intertidal communities. However, we found no significant correlation with air exposure time or fifth percentile air temperatures, despite a transition zone in tidal regime from semidiurnal to mixed tidal which occurs north of Disko Island, and that coincides with decreasing biomasses. Increased exposure time to low air temperatures significantly increases temperature mortality in intertidal benthic species (Aarset, 1982; Thyrring et al., 2017), and although most Arctic intertidal sessile organisms are freeze tolerant (Murphy, 1983; Thyrring et al., 2020a), few species are capable of surviving extended exposure to sub-zero temperatures. Therefore, survival of benthic species on exposed high Arctic rocky shores depends on microhabitats, such as crevices, the ice foot and biogenic habitats, which reduce exposure to extreme sub-zero air temperatures (Scrosati & Eckersley, 2007; Ørberg et al., 2018). So, although we found no direct effects of air temperatures derived from weather stations, extreme low temperature events in Greenland may restrict species to inhabiting protective microhabitats where temperatures deviate significantly from weather station measurements (Helmuth, 1998). For example, a recent study showed that while the average microhabitat temperature on Disko Island was − 3.07°C, temperatures at the nearby weather station was − 8.04°C (Thyrring et al., 2017). Thus, considering temperatures on scales relevant to the organisms will improve predictability of temperature impacts on intertidal assemblage structure and species distribution (Jurgens & Gaylord, 2018).

Although, a clear latitudinal pattern in the duration of seasonal ice coverage, and ice scour is evident, we identified no relation between average annual sea ice coverage or ice scour and intertidal assemblage metrics among the six studied regions. There are several potential explanations for this. One explanation is that the export of ice from northeast to south Greenland with the East Greenland Current may cause higher local ice scour frequencies in south Greenland compared to Nuuk and Disko Island (Høgslund et al., 2014). Also, icebergs are lock up for longer in regions with short ice-free summers, reducing scour from drifting ice (Barnes et al., 2014), and the stress from ice scour depends not only on the seasonal presence of ice, but also on the interaction with wave exposure (Heaven & Scrosati, 2008), which may result in locally low biomasses at wave-exposed sites not captured in the present study. The potential stress from ice-scour also depends on the presence of cracks and crevices, as discussed on temperature stress above. Hence, in suitable microhabitats, large standing stocks can develop in the high Arctic; local biomass of *F. distichus* subsp. *evanescens* was similar in a southern (Nuuk) and northern (Upernavik) region, and 100% within site macroalgal coverage was also found in all six regions where sites were protected. Furthermore, at sheltered sites in Nuuk seasonally exposed to fast ice, the biomass of *A. nodosum* exceeded 20,000 g ww m^-2^, which correspond to biomasses near its distribution center in Canada that range from 11,400 to 28,900 g ww m^-2^ (Vadas et al., 2004). Thus, our results support the notion that high algal biomass and coverage can be found in the Arctic region when local conditions are favorable (Wulff et al., 2009). In summary, the lack of correlation between intertidal assemblage variables and our environment measures partly reflects: 1) the buffering role of small-scale conditions such as the presence of crevices and/or a protective ice foot allowing vegetation to develop despite being covered by fast ice and 2) that drifting ice also cause ice scour.

In the Arctic, vertical zonation patterns have mostly been described at Svalbard where species richness increase from the intertidal towards the subtidal realm (Hop et al., 2002; Wulff et al., 2009). In the present study, we showed that *A. nodosum* is restricted to the low intertidal zone near its northernmost intertidal distribution, and a previous intertidal study showed that blue mussels (*Mytilus* sp.) distribution is progressively restricted downward to the lower stress low intertidal in north Greenland (Thyrring et al., 2017). However, the vertical distribution of the dominant intertidal species shows no change across latitudes; *F. vesiculosus, F. distichus* subsp. *evanescens* and *S. balanoides* consistently occupied >70% of the tidal amplitude. Arctic populations of *Fucus* spp. can survive extended exposure to sub-zero air temperatures without injury (Parker, 1960; Becker et al., 2009), and *M. edulis* and *S. balanoides* can survive air temperatures near −15°C (Cook & Lewis, 1971; Thyrring, Juhl, Holmstrup, Blicher, & Sejr, 2015). Thus, when species utilize protective microhabitats in crevices and cracks, local environmental conditions on scales relevant to the organisms do not induce any vertical changes in the presence of foundation species in north Greenland.

### 4.1 A resilient ecosystem

Ecosystem resilience is defined as a combination of resistance to severe disturbances, capacity for recovery, and the ability to adapt to new conditions (Bernhardt & Leslie 2013). We argue that Greenland’s intertidal assemblage also has an inherent resistance to climate change. The ecosystem is generally dominated by stress tolerance species able to survive extreme environmental stress. This is especially so in microhabitats, which mitigate the impacts of environmental stress, allowing large standing stocks to develop locally across the climate-gradient. It is possible that a future increase in temperature and a reduction in seasonal sea ice coverage will affect intertidal production and thereby standing stock positively, as suggested for *Ascophyllum nodosum* in Greenland (Marbà et al., 2017) and predicted for the subtidal realm (Krause-Jensen & Duarte, 2014; Olesen, Krause-Jensen, Marbà, & Christensen, 2015). The resilience is also reflected in that the most abundant and widely distributed foundation species (i.e. *Fucus* spp.) across the intertidal assemblage, have a strong recovery capacity as they are able to reproduce, grow and recolonize following physical disturbance by ice (Kiirikki & Ruuskanen, 1996; Ørberg et al., 2018). Moreover, recent studies have shown adaptation to local environmental conditions takes place in intertidal Arctic populations of barnacles (*S. balanoides*) (Marshall et al., 2018) and blue mussels (*M. edulis*) (Telesca et al., 2019; Thyrring et al., 2020a). Thus, West Greenland’s intertidal ecosystem seems resilient to gradual climate change, and it is likely that introduced species could pose a greater risk for intertidal assemblages than climate change as increased shipping and propagule transport with the Irminger current increase the potential for introduction of non-native species (Ingolfsson, 1992; Renaud et al., 2015). Most marine ectotherms generally fill their thermal niches (Sunday et al., 2012). Thus, increasing temperatures may allow species currently residing in temperate regions to expand their distribution polewards, and increase their production and biomass in suitable habitats, as observed in other Arctic regions (Węsławski et al., 2010; Kortsch et al., 2012; Fossheim et al., 2015). Predatory species are known to inhabit and structure rocky intertidal communities in northern temperate areas and south Iceland (Ingólfsson, 2004; Jenkins et al., 2008), and an increased inflow of warm Atlantic water to West Greenland in the 1930’s led to large-scale changes in coastal ecosystems, including the arrival of the predatory starfish *Asterias rubens* (Drinkwater, 2006). *Asterias rubens* and the predatory dogwhelk *Nucella lapillus* are found in the sublittoral of West Greenland (e.g. Mortensen 1932; Thorson 1951; Pers. obs MB), and future climate warming may facilitate an expansion of their distribution into the intertidal, likely altering the structural dynamics of assemblages. This study, therefore, serves as an important reference point for potential bottleneck events and for future tests of such predictions, as well as for identification of effects of climate change through future monitoring of the intertidal assemblage in Greenland.

## Acknowledgement

Peter Bondo Christensen and Nuria Marbà are thanked for help with sampling at Nuuk. This work is a contribution to the Arctic Science Partnership (ASP), and to the Greenland Ecosystem Monitoring program (https://g-e-m.dk/).

## Data availability

All raw data necessary to replicate this study is freely available on Zenodo (Link will be added upon acceptance). Weather data is available from the Danish Meteorological Institute (www.research.dmi.dk)

